# Machine learning reveals sequence-function relationships in family 7 glycoside hydrolases

**DOI:** 10.1101/2020.11.06.372003

**Authors:** Japheth E. Gado, Brent E. Harrison, Mats Sandgren, Jerry Ståhlberg, Gregg T. Beckham, Christina M. Payne

## Abstract

Family 7 glycoside hydrolases (GH7) are among the principal enzymes for cellulose degradation in nature and industrially. These important enzymes are often bimodular, comprised of a catalytic domain attached to a carbohydrate binding module (CBM) via a flexible linker, and exhibit a long active site that binds cello-oligomers of up to ten glucosyl moieties. GH7 cellulases consist of two major subtypes: cellobiohydrolases (CBH) and endoglucanases (EG). Despite the critical biological and industrial importance of GH7 enzymes, there remain gaps in our understanding of how GH7 sequence and structure relate to function. Here, we employed machine learning to gain insights into relationships between sequence, structure, and function across the GH7 family. Machine-learning models, using the number of residues in the active-site loops as features, were able discriminate GH7 CBHs and EGs with up to 99% accuracy. The lengths of the A4, B2, B3, and B4 loops were strongly correlated with functional subtype across the GH7 family. Position-specific classification rules were derived such that specific amino acids at 42 different sequence positions predicted the functional subtype with accuracies greater than 87%. A random forest model trained on residues at 19 positions in the catalytic domain predicted the presence of a CBM with 89.5% accuracy. We propose these positions play vital roles in the functional variation of GH7 cellulases. Taken together, our results complement numerous experimental findings and present functional relationships that can be applied when prospecting GH7 cellulases from nature, for sequence annotation, and to understand or manipulate function.

## Introduction

Cellulose is the most abundant renewable biopolymer on Earth and, thus, holds tremendous potential in transitioning energy production from fossil fuels to a renewable carbon feedstock — a key need to limit anthropogenic climate change. Sugars derived from the deconstruction of cellulose can be converted to biofuels and numerous chemicals via myriad biological or catalytic conversion routes. However, the efficient depolymerization of cellulose in a cost-effective manner such that biofuels can economically compete with fossil fuels remains a major challenge to enabling a lignocellulosic economy (1). In industry, biochemical methods of cellulose deconstruction employing enzymes are promising due to high selectivity, low energy consumption, and low amounts of by-product generation (1–3). As a result, improving the yield of enzymatic hydrolysis of cellulose by enhancing cellulase activity is a major research focus.

In nature, microbial cellulose degradation is primarily achieved via a synergistic cocktail of enzymes consisting of processive cellobiohydrolases (CBHs), endoglucanases (EGs), and accessory enzymes such as β-glucosidases and lytic polysaccharide monooxygenases (LPMOs) (2). Organisms can employ these enzymes as free single- or multi-modular constructs, or as cellulosomes. Industry tends to employ free enzyme systems, as filamentous fungal hosts are proficient secretors of these types of cellulose-degrading enzymes. EGs act by attacking internal bonds in cellulose, thus, creating free chain ends. CBHs attach to free chain ends via exo-initiation, or internal regions in the chain via endo-initiation, and processively cleave off cellobiose units as they process along the chain. Cellobiose products are consequently hydrolyzed by β-glucosidases to yield glucose (2). Whereas CBHs are known to be processive and to carry out several cellulolytic cuts before detaching from the cellulose substrate, EGs are mostly nonprocessive or may show little processivity (4–8). Optimum cellulolytic efficiency is achieved by the synergistic action of CBHs and EGs. CBHs, EGs, and β-glucosidases, as well as other glycoside hydrolases (GHs) are currently classified into 168 families in the CAZy database (9,10).

Family 7 glycoside hydrolases (GH7s) are the powerhouses of cellulose degradation in nature. They traditionally are found mostly in fungi, although sequences have been identified in several non-fungal groups such as Crustacea, Porifera, Alveolata, and Amoeba (11). Because GH7s offer significant cellulolytic potential, they are often the predominant enzymes by mass in the secretomes of many filamentous cellulolytic fungi and constitute the major components of enzyme cocktails in industrial cellulolytic processes (2,12,13).

GH7s consist of two main subtypes, CBHs and EGs. Although over 5,000 GH7 sequences are known, structural information is presently available for only 21 GH7s (16 CBHs, 5 EGs) (11,14–31). GH7 CBH and EG structures share a similar β-jelly roll fold with two antiparallel β-sheets that pack into a curved β-sandwich (15). Loops protrude from the β-sandwich and extend over a tunnel-like active site that spans 40–50 Å across the ends of the catalytic domain (CD). The active site contains at least nine glycosyl subsites for binding cello-oligomers, which are numbered −7 to +2 from the non-reducing end of the cellulose chain **(Figure 1)**. The cellulose chain is cleaved between the −1 and +1 subsites (2). Despite the overall similarity in fold, structures of GH7 CBHs and EGs are strikingly different in their activesite configuration. Whereas GH7 CBHs exhibit a closed tunnel-like active site, GH7 EGs possess a more open, groove-like active site. These differences arise due to the variation in the residue lengths of the loops that protrude over the active-site groove, labeled A1 to A4 and B1 to B4 **(Figure 1)** (16).

**FIGURE 1.**
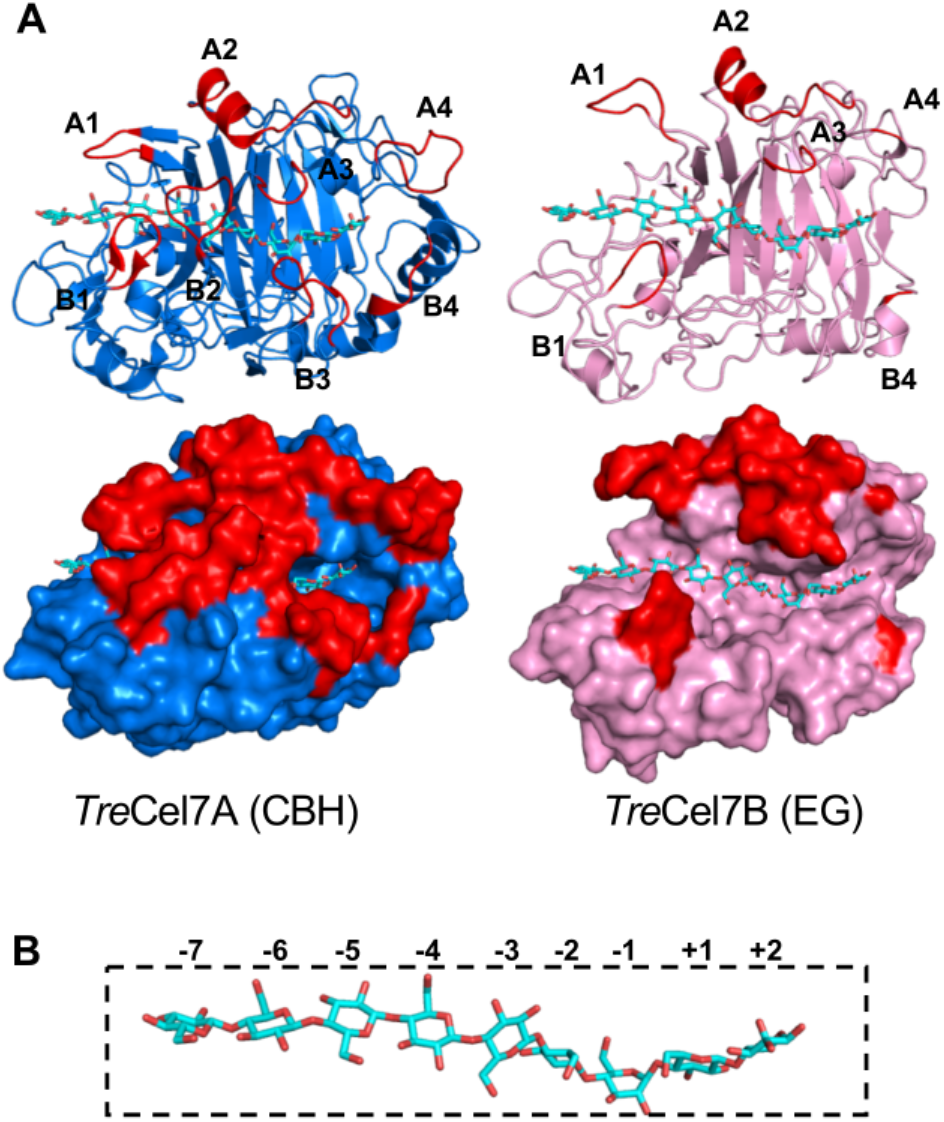
Structures of typical GH7 CBH and EG with a cellononaose ligand in complex. **(A)** The CBH (left), *Trichoderma reesei* Cel7A (*Tre*Cel7A, PDB code: 4C4C) (24), and the EG (right), *Trichoderma reesei* Cel7B (*Tre*Cel7B, PDB code: 1EG1) (27). The eight active-site loops (A1 to A4 and B1 to B4) are shown in red. In the CBH, the active site is tunnel-like, but is more open and groove-like in the EG. **(B)** Glycosyl binding sites are numbered from the non-reducing end at the active-site tunnel entrance (−7) to the reducing end (+2) where the cellobiose product exits the active site. Bond cleavage occurs between −1 and +1 subsites.

Several structural and mechanistic studies of GH7s have proposed that the differences in functional properties of GH7 CBHs and EGs, such as processivity, endo-initiation, and product inhibition, arise mainly due to the differences in the active-site architecture in the loops (16,17,20,22,25,26,29,30). Moreover, GH7 CBHs with a more exposed activesite tend to exhibit functional characteristics intermediate between typical CBH and EG behavior (6,20,32,33). Besides the differences in the configuration of active-site loops, studies have also indicated that there are key residues in the active site of GH7s that contribute to the variation in GH7 CBH and EG behavior. Several aromatic and charged residues in the active site that interact with the cellulose substrate have been suggested to be crucial for the processive activity of GH7 CBHs (23,34–37). Furthermore, mutation of these residues notably diminishes the processive activity of GH7 CBHs on crystalline cellulose (38,39).

Like many other cellulases, GH7s can be bimodular, having their CD attached to a carbohydrate binding module (CBM) by an intrinsically disordered glycosylated linker peptide (40–44). There are currently 87 families of CBMs in the CAZy database (9.45), but GH7s mainly utilize family 1 CBMs (2.46). It is now generally accepted that family 1 CBMs function to increase the affinity of cellulases for crystalline cellulose and, thereby, increase the surface concentration of the enzyme for catalysis. Thus, by facilitating two-dimensional diffusion of the CD on the cellulose surface, the CBM improves catalytic efficiency (40). Furthermore, several studies have revealed that deletion of the CBM-linker domain dramatically reduces CBH activity on crystalline cellulose, especially at low enzyme concentration, but not on soluble substrates (46–51). Takashima *et al*. carried out several mutations in the CBM of a *Humicola grisea* CBH (*HgrCel7A*) and observed high positive correlation between the efficiency of the enzyme on crystalline cellulose and the binding affinity of the CBM (52). Similarly, Srisodsuk *et al*. observed that replacing the CBM of *Trichoderma reesei* Cel7A (*Tre*Cel7A) with the CBM of *Tre*Cel7B, which has a higher cellulose-binding affinity, improved the activity of *Tre*Cel7A on crystalline cellulose (50). Altogether, these results indicate that CBMs affect GH7 catalytic activity primarily by promoting binding to the cellulose surface.

Despite the tremendous growth in scientific knowledge of GH7s over the last few decades, our understanding of how sequence and structure affect function is far from complete. Although it is known that the exposure of the active site due to truncation in the active-site loops can substantially affect function, little work has been done to elucidate the unique roles that each of the active site loops play and how the effects of truncation vary with function for the different loops. Recently, Schiano-di-Cola *et al*. studied the effects of deletions in the B2, B3, and B4 loops on the activity and kinetics of *Tre*Cel7A (53). They found that deletions in the B2 loop, compared to the B3 and B4 loop, most significantly affect CBH behavior of *Tre*Cel7A. Beyond *Tre*Cel7A, there is a need to investigate how variation of active-site loop lengths relate to function across other members of the GH7 family.

In this work, we employ machine learning (ML) to derive relationships between sequence, structure, and function of GH7s using a dataset of 1,748 selected protein sequences. The sequences are aligned via multiple sequence alignment (MSA) to identify regions of structural similarity and evolutionary importance. Although manual inspection of the MSA may reveal several functional patterns, such as highly conserved positions, many important but complex relationships may be missed. ML is an especially useful statistical tool when data are abundant and relationships in the data are complex (54). Thus, ML can be employed to discover complex functional and evolutionary relationships in proteins. In this work, we apply ML to the MSA of GH7 sequences, mapping variation in lengths of the active-site loops to functional subtypes such that the subtype can be accurately predicted from loop length. We also derive position-specific classification rules to highlight positions that play important roles in CBH/EG function. Lastly, we investigate relationships between the CBM and the CD by utilizing ML to predict the presence of CBMs in GH7s using residues in the CD. It is important to note that, as the current understanding of GH7 function is based on investigation of a few representatives, this present study of 1,748 GH7 sequences seeks to identify general sequence-function relationships for the entirety of the GH7 family and the degree to which variation exists.

## Results

### Datasets

Three datasets were used in this study. The first dataset contained 1,748 full-length GH7 protein sequences retrieved from the National Center for Biotechnology Information (NCBI) non-redundant database. Using a strict keyword search, we queried the NCBI database for the subtype annotation (i.e. CBH or EG) of these 1,748 sequences. 427 sequences were clearly annotated as CBH or EG in the database (291 CBHs and 136 EGs), and these 427 sequences comprised the second dataset. For the third dataset, we retrieved 44 GH7 sequences from the manually curated UniProtKB/Swiss-Prot database (55). Accordingly, the subtype annotations of the 44 GH7s (30 CBHs, 14 EGs) are less likely to contain errors than the annotations of the 427 sequences from the NCBI non-redundant database.

### Discrimination of GH7 subtypes with hidden Markov models

In the annotation of a protein sequence, several computational prediction methods may be applied. Sequence similarity methods compare an unclassified protein with well-studied proteins and assign the unclassified protein to the same class as the most similar classified proteins (56). Hidden Markov model (HMM) (57,58), which describes the protein sequence as a probabilistic model, is one of the most sensitive and most accurate methods for discriminating protein functional families with sequence data alone, provided they are built with correct alignments (56). Within a given protein family, HMM can also be applied to discriminate functional subtypes, although the discrimination accuracy varies across different families (59).

We applied HMM to discriminate GH7 CBHs and EGs. The performance of HMM was evaluated by a five-fold cross-validation technique using the datasets of 427 (NCBI) and 44 (UniProtKB/Swiss-Prot) GH7 sequences. First, each dataset was aligned and separated into CBH and EG subalignments based on the database annotations. Then, each subalignment was randomly split into five folds **(Figure 2A)**. Subtype HMMs (i.e., CBH HMM and EG HMM) were repeatedly built on four out of five folds of the CBH and EG subalignment, and the sequences in each left-out fold were used as a test set. To predict the subtype of a sequence, the sequence was aligned separately to both the CBH and EG HMMs, and then the alignment scores were compared. If the CBH HMM alignment score was greater than the EG HMM alignment score, the sequence was predicted to be a CBH; otherwise, it was predicted to be an EG (59). The process was repeated so that all five folds were used in training and testing the HMMs.

**FIGURE 2.**
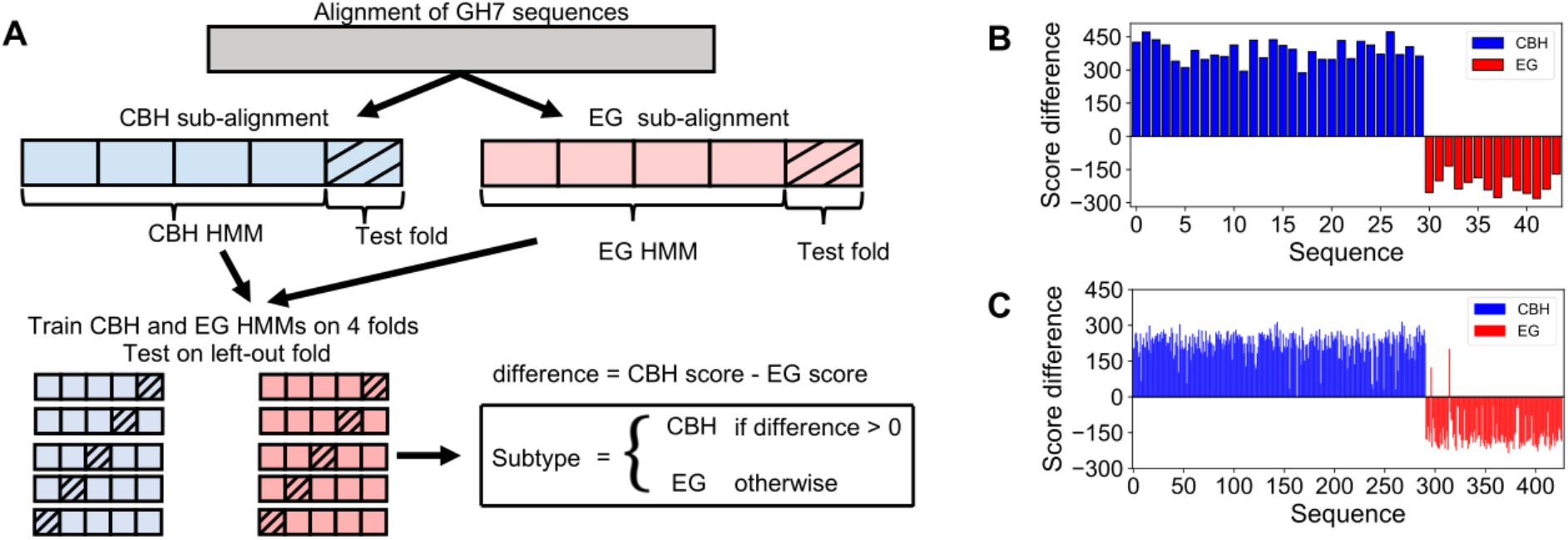
Discrimination of GH7 CBHs and EGs with hidden Markov models (HMM). **(A)** Five-fold crossvalidation technique for evaluating the performance of HMM. The MSA is split into CBH and EG subalignments and each subalignment into five folds. HMMs are repeatedly trained on four folds and then tested on the left-out fold. The predicted class (CBH or EG) of a sequence is the class that yields the highest HMM alignment score. **(B)** Performance of HMM on the dataset of 44 GH7s from the manually-curated UniProtKB/SwissProt database. **(C)** Performance of HMM on the dataset of 427 GH7s from NCBI non-redundant database. Only two EG sequences (GenBank accession codes: AGY80096.1 and AGY80097.1) were misclassified in the NCBI dataset. Note that in B and C, the assigned sequence numbers (*x*-axes) are arbitrary.

**Figure 2B and C** show the performance of the HMM method on the UniProtKB/Swiss-Prot dataset (44 sequences) and on the NCBI dataset (427 sequences), respectively. The HMM method achieved perfect accuracy on the UniProtKB-Swiss-Prot dataset. All sequences were correctly predicted, and there was a substantial difference, of at least 120.0, between the CBH alignment score and the EG alignment score. On the NCBI dataset of 427 sequences, which may contain erroneous subtype annotations, the HMM achieved an accuracy of 99.53% and only misclassified two sequences (accession codes: AGY80096.1 and AGY80097.1), which are annotated as EGs. These two sequences may have been erroneously annotated as EGs since they are much more similar to CBHs in overall sequence and loop lengths. Furthermore, the value of the alignment score difference for some sequences in the NCBI dataset is as low as 2.0.

### Discrimination of GH7 subtypes with machine learning: relationships between active-site loops and CBH/EG function

In this part of the study, our goal was to use ML to map the variation in amino acid sequence to GH7 CBH and EG activity and to, consequently, determine which aspects of the sequence and structure predominantly affect CBH/EG function. If a particular feature is important for the difference in CBH and EG behavior, we should be able to train ML models on that feature to discriminate GH7 CBHs and EGs with significant accuracy. Otherwise, a feature that has no correlation with activity, but only varies due to phylogenetic diversity, would perform poorly when applied to predict GH7 subtypes with ML.

We used the dataset of 1,748 GH7s to test ML algorithms in predicting GH7 subtypes. Since only 427 of the 1,748 GH7s are classified as CBH or EG in the databases, we applied the HMM method described previously to derive the functional classes of the unclassified GH7 sequences. Our crossvalidation tests showed that the HMM method can correctly classify GH7 subtypes with an accuracy of almost 100% (i.e. consistent with the database annotations). This result is similar to the performance of the HMM method applied to other protein families (59). Moreover, when we trained separate HMMs on the manually-annotated dataset of 44 sequences (UniProtKB/Swiss-Prot) and on the “less perfect” dataset of 427 sequences (NCBI), and then applied the HMMs to determine the subtype of the 1,748 GH7s, the separate HMMs assigned the same subtype in all but five instances (99.71%). Regardless, misclassification errors of about 1% are not large enough to alter the relationships that we derived from ML on the dataset of 1,748 GH7 sequences (60,61).

In choosing features for the ML models, we capitalized on the observation that crystal structures of GH7 CBHs and EGs differ in their active-site architecture, due to the degree of truncation in the eight active-site loops **(Figure 1).** Hence, we used the number of residues in the active-site loops as features for ML to discriminate between GH7 CBHs and EGs. First, a structure-based MSA of all 1,748 sequences was carried out (See Materials and Methods for details). For each sequence in the MSA, we counted the number of amino acid residues in the eight active-site loops and derived a vector of the eight loop lengths as features **(Figure 3)**.

**FIGURE 3.**
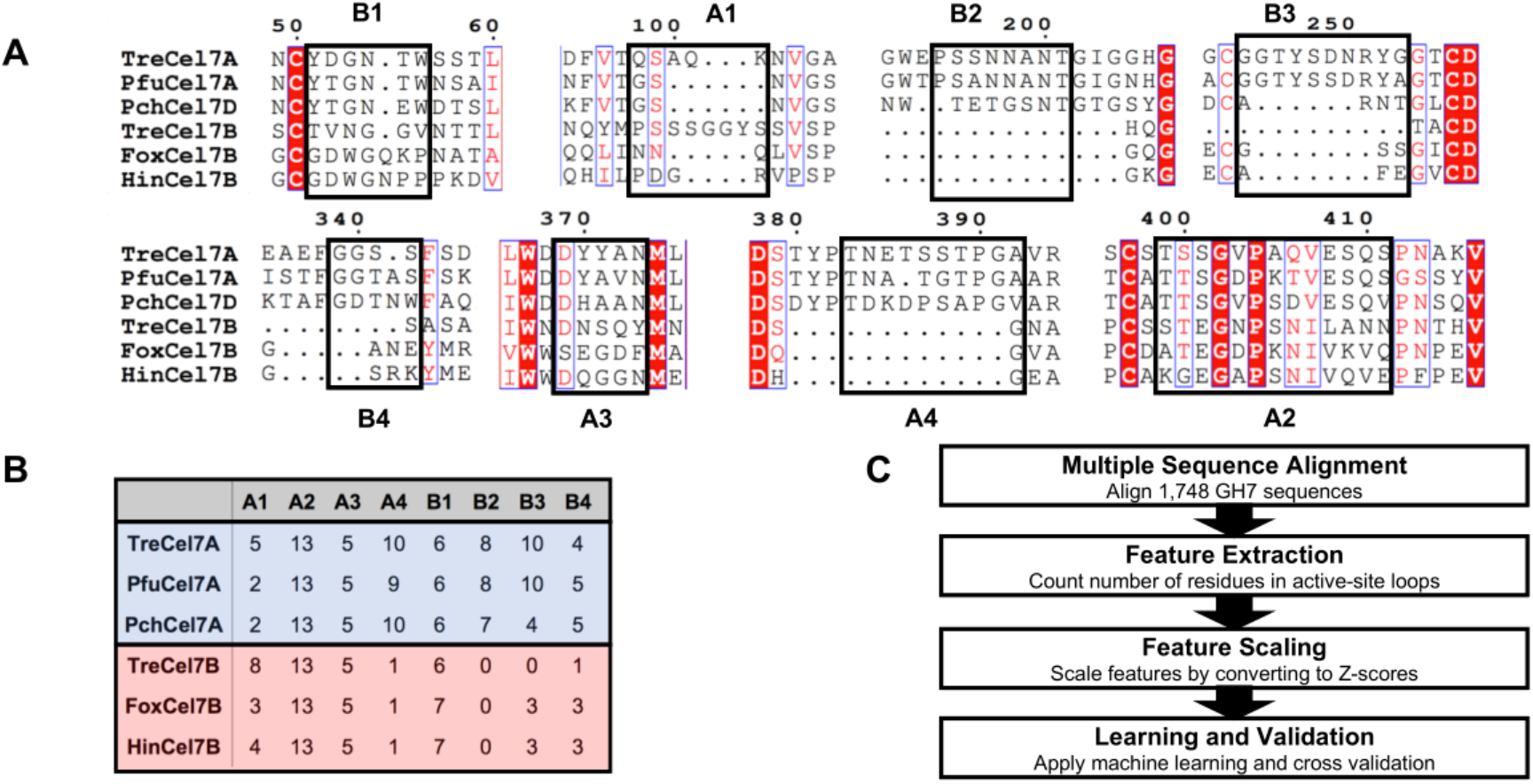
Generating features for discriminating GH7 CBHs and EGs with ML. **(A)** Segments of a selection of six well-studied GH7s from the structure-based sequence alignment of 1,748 sequences showing the activesite loops. The sequences include the CBHs: *Trichoderma reesei* Cel7A (*Tre*Cel7A) (24), *Penicillium funiculosum* Cel7A (*PfuCel7A*) (21), and *Phanerochaete chrysosporium* Cel7D (*Pch*Cel7D) (20); and the EGs: Trichoderma reesei Cel7B (*Tre*Cel7B) (27), *Fusarium oxysporum* Cel7B (*Fox*Cel7B) (25), and *Humicola insolens* Cel7B (*Hin*Cel7B) (26). **(B)** The number of residues in the eight active-site loops as determined from the structure-based alignment. **(C)** Procedure for generating features for 1,748 GH7s. First, the sequences are aligned as in (A). Then, a count of the number of residues in each loop is obtained. Residue counts are scaled to Z-scores before ML is applied.

Four ML methods were applied: decision trees, logistic regression, k-nearest neighbors (KNN), and support vector machines (SVM). For each ML method, nine models with different combinations of features were tested. One model involved training the ML algorithms on the lengths of all eight loops, and the remaining 8 models involved using each loop length as the sole feature for the training (single-feature models). The performance of the ML models was measured using four metrics: sensitivity (or true positive rate), specificity (or true negative rate), overall accuracy, and Matthew’s correlation coefficient (MCC). Here, the sensitivity is the percent of CBHs (the true class) correctly predicted, the specificity is the percent of EGs (the false class) correctly predicted, and the overall accuracy is the percent of both CBHs and EGs correctly predicted. MCC ranges from −1 to +1 and measures the correlation between the predicted and true classifications. An MCC value of +1 indicates perfect prediction, 0 indicates no concordance between predicted and actual classes, and −1 indicates perfect disagreement. MCC has been recommended as the most informative performance metric in evaluating binary classification performance, especially when the dataset is imbalanced since other metrics such as overall accuracy and F1 score can be hugely misleading (62–65). Hence, we use MCC as the primary metric in evaluating the performance of the ML models.

Moreover, we are faced with the problem of an imbalanced dataset: 1,306 (75%) of the 1,748 sequences in the dataset are CBHs. Ordinarily, imbalanced data will skew the results by causing the ML classifiers to place most of the data in the majority class (CBH). To deal with the imbalance problem, we applied random undersampling (66,67) to the majority class so that the distribution of CBH and EGs was balanced. We evaluated the performance of the ML models on the redistributed data with 100 repetitions of five-fold cross validation, with the dataset undersampled and reshuffled in each repetition **(Figure 4)**. Repeating the five-fold cross validation numerous times is a highly effective way to mitigate the effects of variability in the train-test splits and to ensure that the data space is thoroughly explored despite loss of data in the undersampling step (68).

**FIGURE 4.**
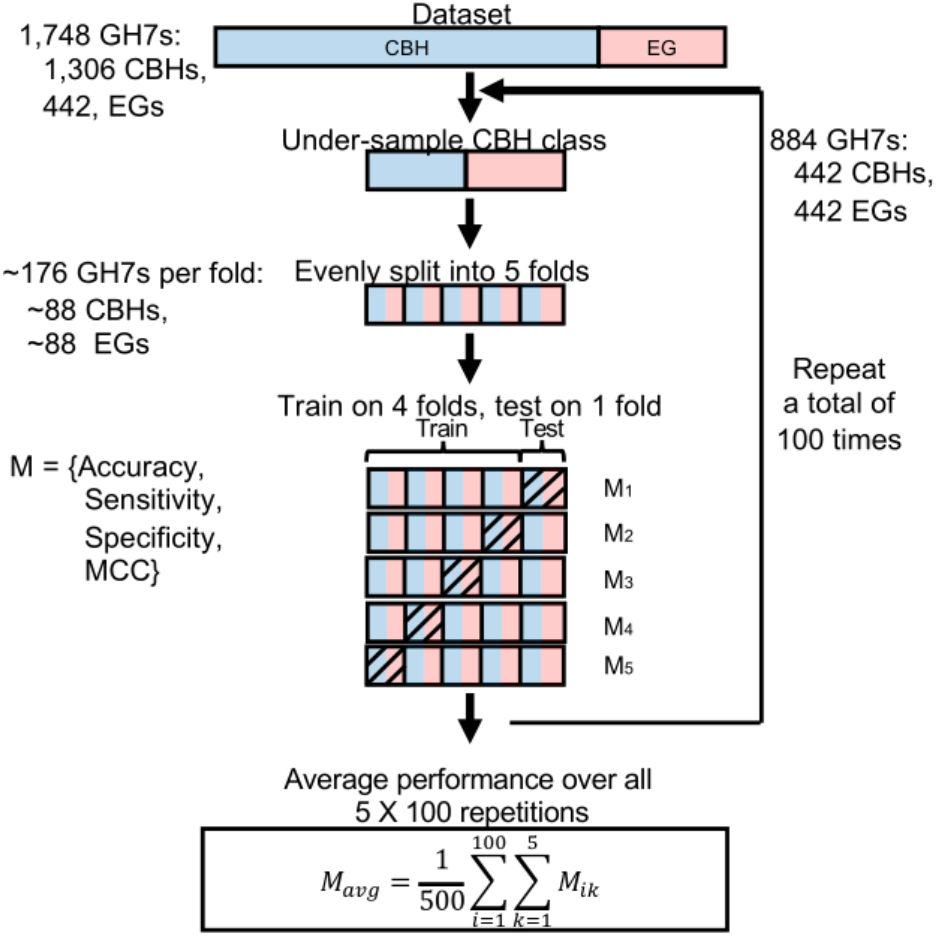
Procedure for evaluating the performance of ML models using 100 repetitions of five-fold cross validation with undersampling. The dataset is reshuffled and resampled in each repetition.

Our results show that ML is able to accurately discriminate between GH7 CBHs and EGs using only information about the length of the active-site loops **(Table 1)**. However, the performance varied significantly for the different single-feature models **(Figure 5A)**. The models trained on the A2 and A3 loops exhibited the worst performance with MCC values close to zero, indicating that they did not perform better than a random classification. The models trained on A1 and B1 loops showed intermediate performance with MCC values widely varying from −0.08 to 0.79 for the A1 models, and −0.03 to 0.63 for the B1 models. Interestingly, the A4, B2, B3, and B4 models showed very high predictive performance, with MCC values ranging from 0.94 to 0.98 and with much lower variation among the different ML methods. The models trained on these five loops (A4, B2, B3, B4) achieved nearly the same high performance as the models trained on all eight loops **(Table 1).**

**Table 1.** Performance of ML algorithms in discriminating GH7 CBHs and EGs*.

|  | Decision tree |  |  | Logistic regression |  |  | K-nearest neighbor |  |  | Support vector machine |  |  |
| --- | --- | --- | --- | --- | --- | --- | --- | --- | --- | --- | --- | --- |
| Features | Sens. | Spec. | Acc. | Sens. | Spec. | Acc. | Sens. | Spec. | Acc. | Sens. | Spec. | Acc. |
| A1 | 98.6 ± 1.2 | 45.9 ± 5.0 | 72.3 ± 3.2 | 42.0 ± 16.5 | 52.8 ± 7.3 | 46.9 ± 6.4 | 86.5 ± 15.1 | 88.7 ± 5.4 | 87.6 ± 5.6 | 97.0 ± 1.8 | 85.5 ± 3.4 | 91.2 ± 2.0 |
| A2 | 65.9 ± 43.5 | 37.4 ± 42.2 | 50.7 ± 3.7 | 49.3 ± 46.7 | 50.3 ± 45.3 | 47.4 ± 2.8 | 4.6 ± 2.3 | 97.0 ± 1.9 | 50.8 ± 3.8 | 89.2 ± 27.3 | 18.4 ± 26.0 | 53.0 ± 3.5 |
| A3 | 89.0 ± 26.3 | 16.9 ± 24.8 | 52.5 ± 3.7 | 50.8 ± 47.9 | 49.4 ± 45.4 | 47.6 ± 3.2 | 3.0 ± 2.0 | 97.8 ± 1.6 | 50.4 ± 3.7 | 96.7 ± 11.2 | 11.4 ± 10.9 | 53.9 ± 3.4 |
| A4 | 95.7 ± 2.1 | 99.5 ± 0.7 | 97.6 ± 1.1 | 95.8 ± 2.0 | 99.5 ± 0.6 | 97.7 ± 1.1 | 95.8 ± 2.1 | 99.7 ± 0.5 | 97.8 ± 1.1 | 95.6 ± 2.2 | 99.6 ± 0.6 | 97.6 ± 1.1 |
| B1 | 96.8 ± 1.8 | 44.1 ± 5.5 | 70.5 ± 3.3 | 79.3 ± 35.6 | 34.5 ± 12.2 | 55.9 ± 13.4 | 1.3 ± 1.8 | 98.7 ± 1.6 | 50.0 ± 3.7 | 95.1 ± 2.6 | 72.3 ± 4.4 | 83.7 ± 2.6 |
| B2 | 94.6 ± 2.4 | 99.1 ± 1.2 | 96.9 ± 1.3 | 94.7 ± 2.4 | 98.4 ± 1.2 | 96.6 ± 1.3 | 95.3 ± 2.3 | 97.4 ± 1.8 | 96.4 ± 1.3 | 94.8 ± 2.4 | 98.4 ± 1.6 | 96.6 ± 1.4 |
| B3 | 92.4 ± 2.7 | 99.8 ± 0.5 | 96.1 ± 1.4 | 89.9 ± 3.3 | 99.8 ± 0.5 | 94.8 ± 1.7 | 96.3 ± 2.2 | 98.6 ± 1.1 | 97.5 ± 1.2 | 89.7 ± 3.3 | 99.8 ± 0.4 | 94.8 ± 1.7 |
| B4 | 97.9 ± 1.8 | 98.2 ± 1.3 | 98.0 ± 1.0 | 98.2 ± 1.4 | 98.2 ± 1.2 | 98.2 ± 0.9 | 97.6 ± 2.0 | 98.3 ± 1.3 | 98.0 ± 1.1 | 97.8 ± 1.6 | 98.2 ± 1.3 | 98.0 ± 1.0 |
| All 8 loops | 98.8 ± 1.2 | 99.1 ± 1.1 | 98.9 ± 0.8 | 98.3 ± 1.4 | 99.2 ± 0.9 | 98.8 ± 0.8 | 97.1 ± 2.0 | 99.4 ± 0.7 | 98.2 ± 1.1 | 99.0 ± 1.1 | 98.9 ± 1.1 | 98.9 ± 0.7 |

\*Each ML model was separately trained separately with each of the eight loops as a single, independent feature, and then with all eight loops combined (last row). The performance was evaluated by measuring the sensitivity (sens.), specificity (spec.) and accuracy (acc.) in percent. Error represents one standard deviation from the mean.

**FIGURE 5:**
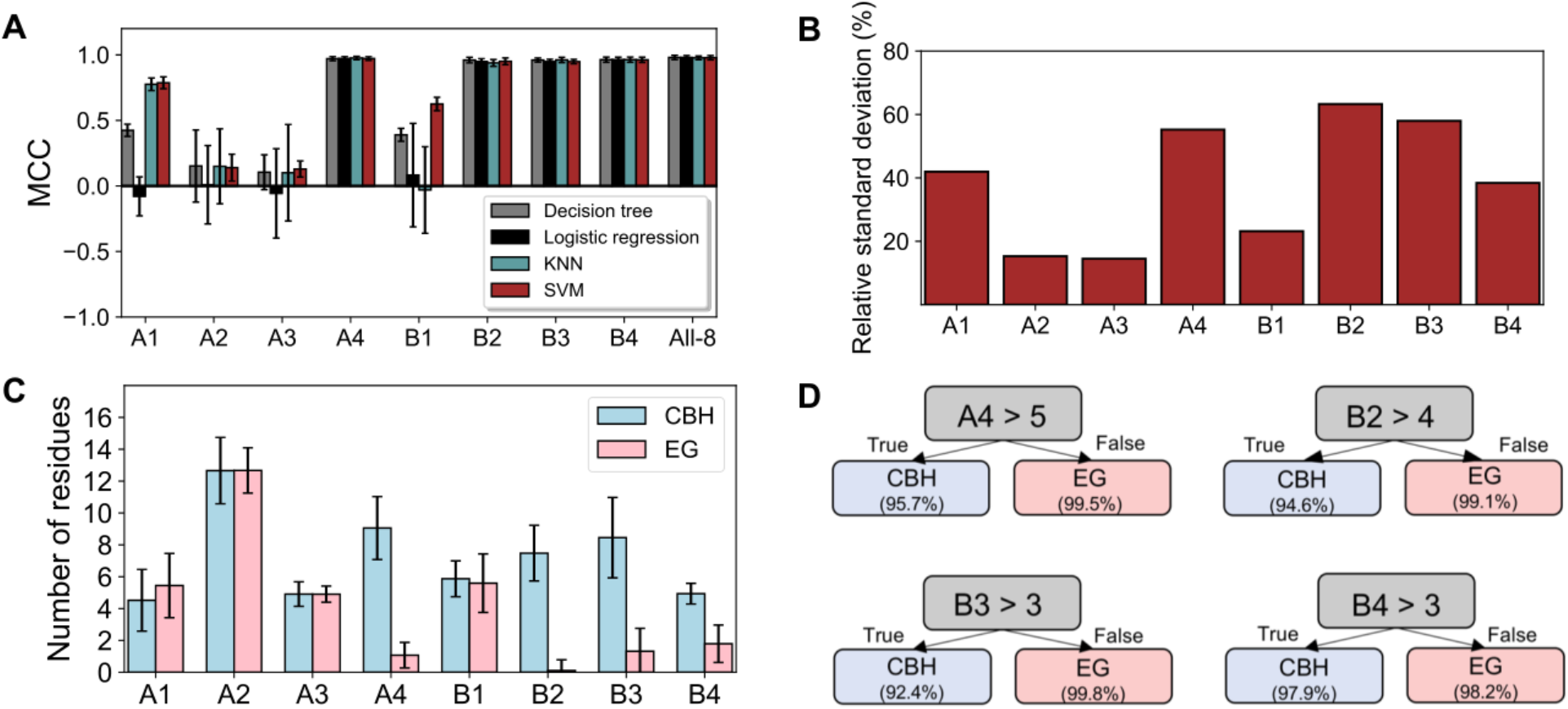
Predictive performance and variation of active-site loops in GH7s **(A)** Matthews’ correlation coefficient (MCC) values of four ML algorithms trained separately on the length of each active-site loop and on all eight loops together. The A4, B2, B3, and B4 loops achieve near-perfect performance in discriminating 1,748 GH7 CBHs and EGs. **(B)** The relative standard deviation of the length of the eight active site loops. Generally, variation in the length of a loop correlates with predictive performance of the loop as a ML feature. **(C)** The mean length of active-site loops in 1,306 GH7 CBHs and 442 GH7 EGs. Error bars are ± 1 standard deviation. **(D)** Rules derived from the single-node decision trees trained on the A4, B2, B3, and B4 loops. The accuracy of the rules in discriminating GH7 CBHs and EGs, i.e. the sensitivity and specificity, respectively, are shown in brackets.

Furthermore, we observed that the variation in the lengths of the loops correlates with the discriminative performance of the loops **(Figure 5A-C)**. The loops with very poor discriminatory performance (A2 and A3) show the lowest relative variation in lengths across the 1,748 GH7s, and nearly identical distributions between CBHs and EGs **(Fig. S1)**. In contrast, loops with intermediate discriminatory performance (A1 and B1) show a greater level of variation in lengths than A2 and A3 loops and noticeably different distributions for CBHs and EGs, although there is a considerable amount of overlap. The loops with near-perfect predictive performance (A4, B2, B3, B4) show the highest variation in lengths.

One major advantage of the tree-based methods over other ML algorithms is the possibility of deriving and visualizing interpretable classification rules (69,70). In many applications of ML to biological problems, it is desirable to gain knowledge of biological relationships rather than merely apply ML as a predictive tool. **Figure 5D** shows rules derived from the single-node decision-tree classifiers trained on the A4, B2, B3, and B4 loops. A classification accuracy of 96.9% was achieved by the simple rule: if a GH7 has more than four residues in the B2 loop, then it is a CBH, else it is an EG. Overall, the decision trees reveal that GH7 EGs tend to possess three or less residues in the B3 and B4 loops, four or less residues in the B2 loop, and five or less residues in the A4 loop.

Since the lengths of the A4, B2, B3, and B4 loops can independently discriminate between GH7 CBHs and EGs with accuracies greater than 94%, it is expected that there is a substantial degree of correlation between them. We conducted correlation analysis by computing the Pearson’s correlation coefficient between the lengths of the eight loops of 1,748 GH7s **(Figure 6)**. As expected, there is significant positive correlation between the lengths of the A4, B2, B3, and B4 loops (r ≥ +0.76, *p* < 0.0001). The highest correlations are observed between the A4 and B2 loops (+0.84) and between the A4 and B4 loops (+0.83).

**FIGURE 6:**
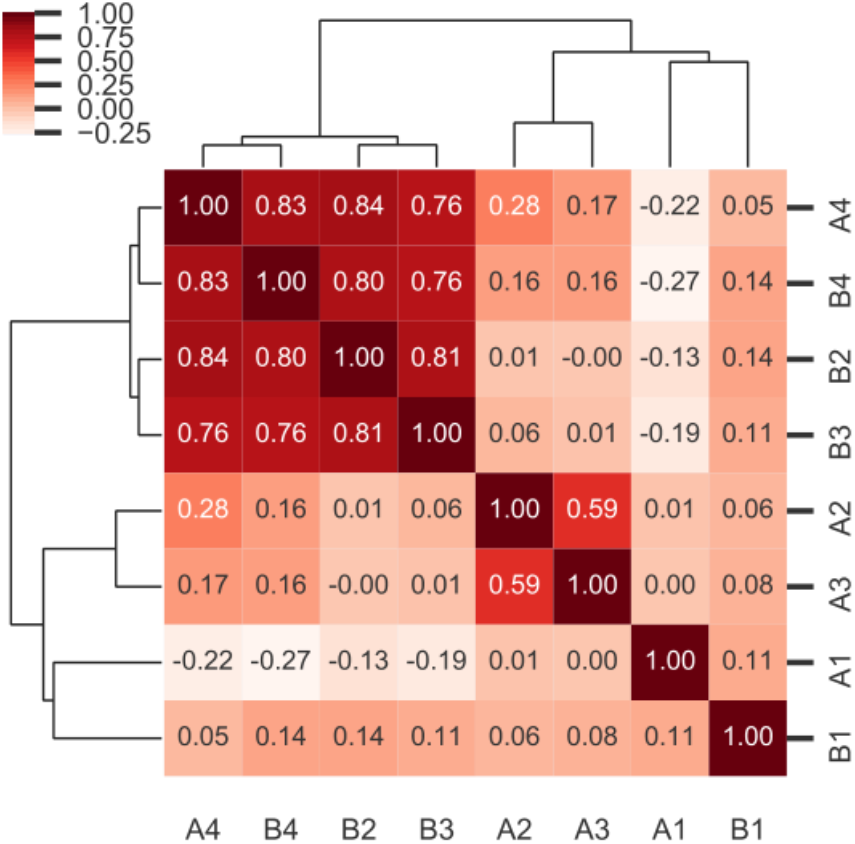
Pearson’s correlation coefficient between the lengths of the eight active-site loops in 1,748 GH7s. The matrix of correlation coefficients is clustered so that loops with a similar pattern of correlation are grouped together. There is a high degree of positive correlation (darker red) between the lengths of the A4, B2, B3, and B4 loops.

### Discrimination of GH7 subtypes with position-specific classification rules: important residues for CBH/EG function

In discriminating GH7 CBHs and EGs with ML, we have used only the lengths of the active-site loops as features without considering the contributions of specific amino acids in the proteins. However, the interactions of specific residues are known to affect GH7 CBH/EG function, and mutagenesis studies have confirmed that certain positions play essential roles in GH7 activity (6,38,39,71). In this section, we investigate the relationships between specific residues in the proteins and the functional subtype.

It is common knowledge that although a protein’s function arises from the combined effects of multilevel interactions between all residues in the protein, some residues contribute to function more significantly than others. Consequently, it is likely that in GH7s, if a position is considerably conserved in CBHs such that CBHs tend to utilize a particular amino acid at that position, and EGs tend to not utilize the same amino acid at that position, or vice versa, then that position plays a vital role in the difference in CBH/EG function or structural stability. A typical example is position 40 (i.e. Trp40 in *Tre*Cel7A). From analysis of the structure-based MSA, we observe that this position is strongly conserved in CBHs with 92.5% exhibiting a Trp at this position, whereas it is notably variable in EGs with only 28.5% exhibiting a Trp at this position **(Figure 7A)**. Considering only this clear difference in the amino acid distribution at position 40, we can infer that Trp40 likely contributes to CBH function. Mutation of Trp40 to Ala has, in fact, been shown to considerably decrease the activity of *Tre*Cel7A on crystalline cellulose but not on amorphous cellulose (38), indicating that Trp40 is critical for processivity (39). Consequently, we propose that applying a statistical method to mine for positions in GH7s that are conserved but have remarkably different amino acid distributions between CBHs and EGs can identify positions that play critical roles in CBH/EG function and processivity.

**FIGURE 7:**
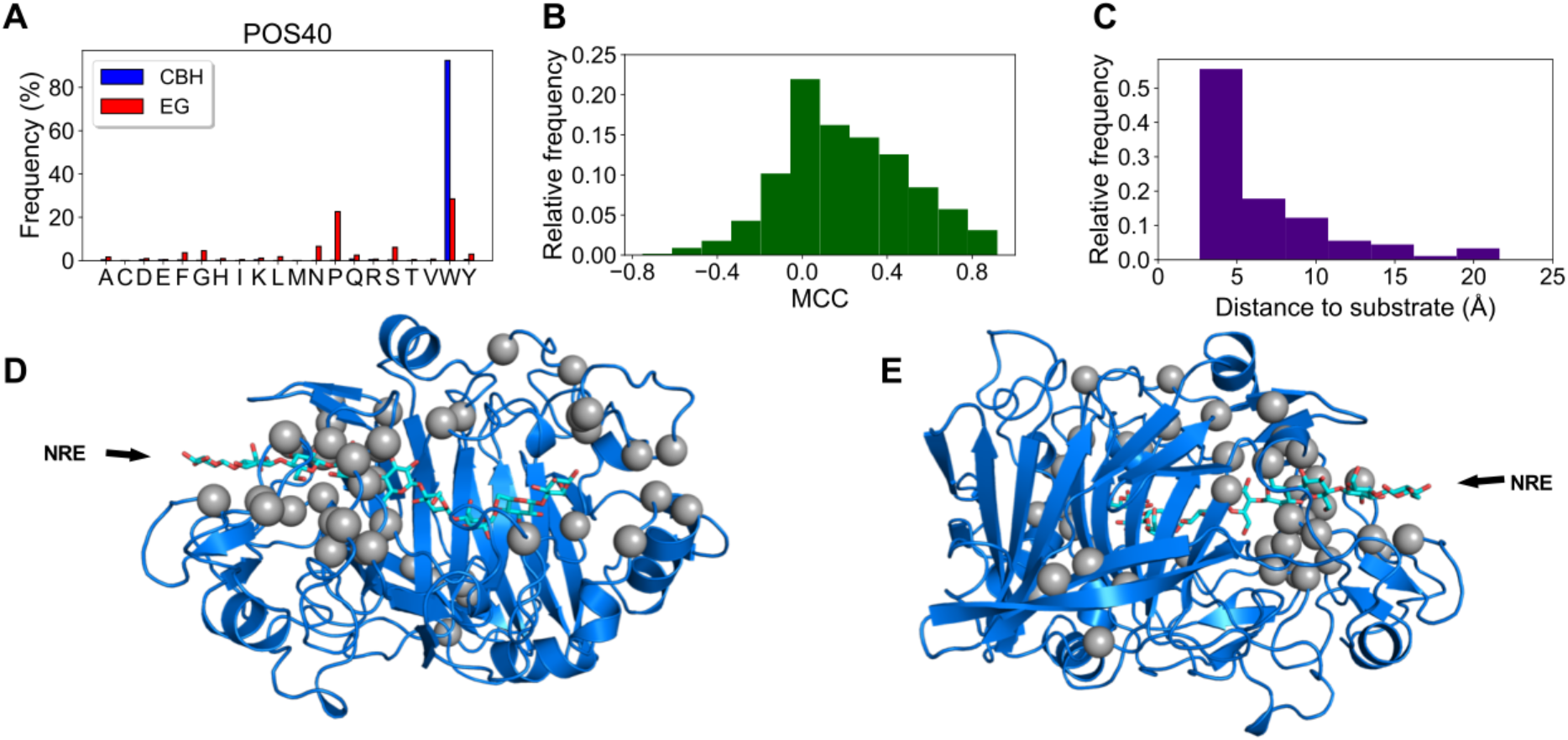
Top-performing position-specific classification rules for discriminating GH7 CBHs and EGs. **(A)** Amino acid distribution of GH7 CBHs and EGs at position 40 (*Tre*Cel7A numbering). Position 40 is strongly conserved as Trp in GH7 CBHs but not in EGs. **(B)** MCC scores of 1,799 position-specific classification rules derived from the MSA. The top 90 rules have MCC scores of 0.73 or greater. **(C)** Histogram of minimum distance between the cellononaose ligand in *Tre*Cel7A (PDB code: 4C4C) (24) and positions from which the top 90 classification rules are derived. More than half of top 90 rules are derived from positions within 5 Å of the substrate. **(D)** Alpha carbons of 42 positions from which the top 90 classification rules are derived shown on the structure of *Tre*Cel7A. Most of these positions are near the substrate sites towards to the nonreducing end (NRE). (E) Posterior view of crystal structure.

From the amino acid distribution at position 40, we obtain a single-node decision tree with the rule: Trp at position 40 implies CBH, else EG. This simple rule classifies 1,748 GH7 CBHs and EGs with an accuracy of 87.2%. Thus, a rational strategy for identifying positions likely associated with CBH/EG function is to derive similar rules for all positions in the MSA and select positions which yield high-performing rules. First, we split the MSA of 1,748 GH7 sequences into CBH and EG subalignments and then identified the consensus amino acid and the consensus amino acid type (i.e. aliphatic, aromatic, polar, positive, or negative) for each position in the subalignments. For each position, if X and Z are the consensus amino acids (or type) in the CBH and EG subalignment, respectively, we derived the following classification rules: X=>CBH and Z=>EG, X=>CBH and not X=>EG, and not Z=>CBH and Z=>EG. Applying this strategy to 434 positions in the MSA (*Tre*Cel7A numbering), we derived 1,799 classification rules. For each rule, we measured the classification accuracy, sensitivity, specificity, and MCC, and tested the statistical significance by conducting chi-square test of independence. The 1,799 rules fairly have normally distributed MCC scores (**Figure 7B**), and the top five percent of rules (90 rules) have MCC scores of at least +0.73, and classification accuracies of at least 87% **(Table 2 and S1, Figure 7 and S2)**. These 90 rules are derived from 42 positions which are generally in close proximity to the cellodextrin ligand in the crystal structure. More than half of the top 90 rules are from positions within 5 Å of the cellononaose ligand bound in *Tre*Cel7A structure (PDB code: 4C4C). Moreover, most of the positions are closer to the tunnel entrance where cellulose chains are recruited by the enzyme for processive hydrolysis **(Figure 7D and 7E)**.

**Table 2.**
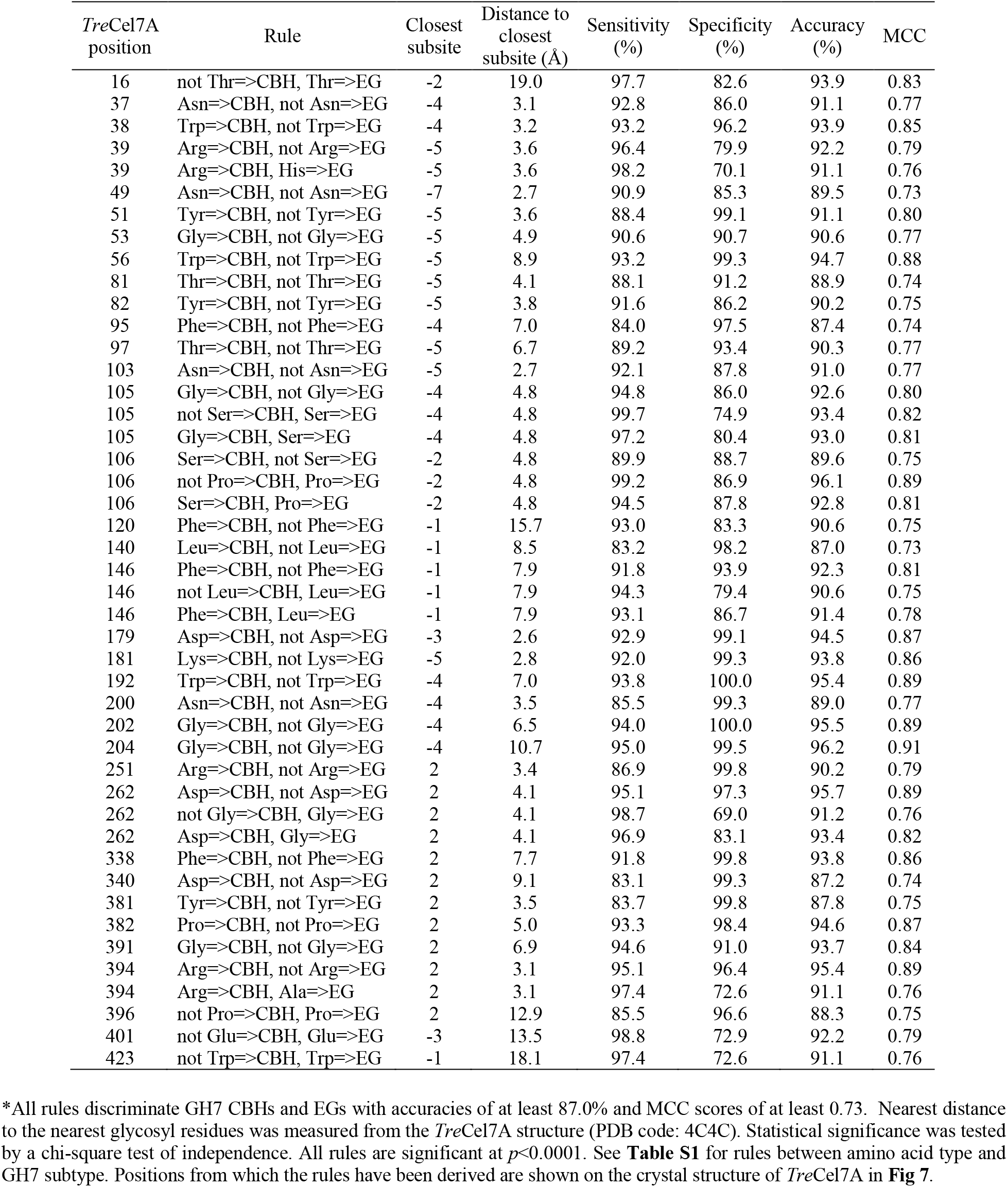
Top-performing position-specific classification rules relating amino acid residues and GH7 subtype (CBH/EG)*.

### Conserved aromatic residues in the active site of GH7s

GH7s possess several aromatic residues lining the active-site tunnel which have been suggested to play key roles in cellulolytic bond cleavage and processive action (36). We have conducted bioinformatic analysis of conserved aromatic residues in the active site of GH7s. From the MSA of 1,748 GH7s, we selected positions that are located within 6 Å of the cellononaose substrate in the structure of *Tre*Cel7A (PDB code: 4C4C), and that have aromatic residues (Phe, Trp, Tyr, or His) at that position in the consensus sequence of CBHs or EGs **(Figure S3)**. There are 17 of such aromatic positions in the MSA, and on the protein structure, these positions are distributed across the nine glycosyl subsites.

Furthermore, these 17 positions can be classified into three groups based on the conservation of aromatic amino acids **(Table 3)**. The first group consists of positions that are conserved in both CBHs and EGs such that more than two-thirds of CBHs and EGs utilize aromatic residues at these positions. Positions 145, 171, 216, 228, 367, and 376 (*Tre*Cel7A numbering) fall in the first group. The second group consists of positions that are conserved as aromatic residues (>66%) in CBHs but not in EGs. Positions 38, 40, 51, 82, 252, 370, and 381 fall in the second group. The third group contains positions that are neither conserved (<66%) as aromatic residues in CBHs and EGs although the consensus amino acids are aromatic. Positions 39, 47, 53, and 247 fall in the third group.

**Table 3.**
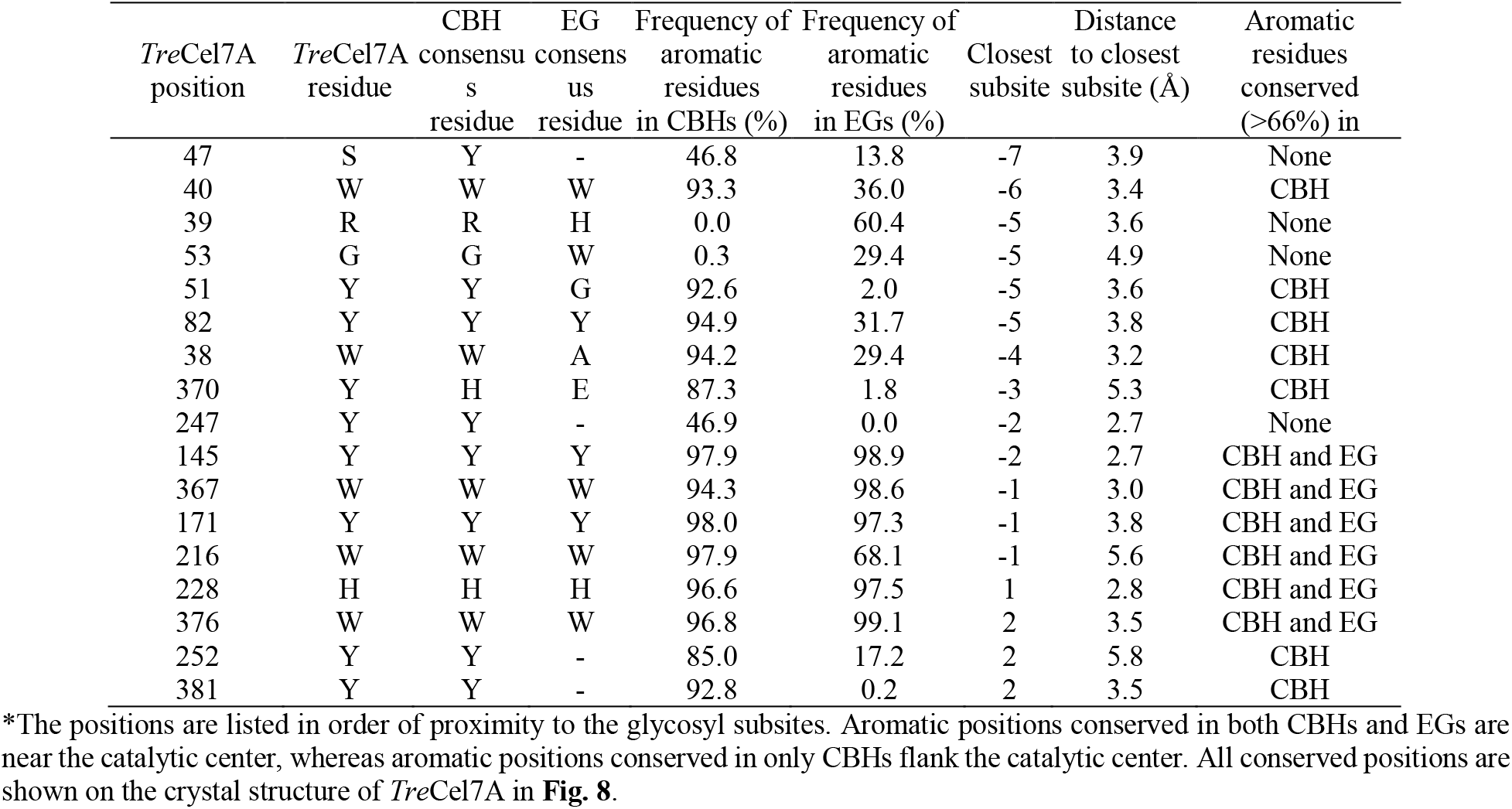
Positions within 6 Å of the cellononaose ligand in *Tre*Cel7A (PDB code: 4C4C) with aromatic residues in CBH/EG consensus sequences*.

| *TreCel7A*<br>position | *TreCel7A*<br>residue | CBH<br>consensus<br>residue | EG<br>consensus<br>residue | Frequency of<br>aromatic<br>residues<br>in CBHs (%) | Frequency of<br>aromatic<br>residues<br>in EGs (%) | Closest<br>subsite | Distance<br>to closest<br>subsite (Å) | Aromatic<br>residues<br>conserved<br>(>66%) in |
| --- | --- | --- | --- | --- | --- | --- | --- | --- |
| 47 | S | Y | - | 46.8 | 13.8 | -7 | 3.9 | None |
| 40 | W | W | W | 93.3 | 36.0 | -6 | 3.4 | CBH |
| 39 | R | R | H | 0.0 | 60.4 | -5 | 3.6 | None |
| 53 | G | G | W | 0.3 | 29.4 | -5 | 4.9 | None |
| 51 | Y | Y | G | 92.6 | 2.0 | -5 | 3.6 | CBH |
| 82 | Y | Y | Y | 94.9 | 31.7 | -5 | 3.8 | CBH |
| 38 | W | W | A | 94.2 | 29.4 | -4 | 3.2 | CBH |
| 370 | Y | H | E | 87.3 | 1.8 | -3 | 5.3 | CBH |
| 247 | Y | Y | - | 46.9 | 0.0 | -2 | 2.7 | None |
| 145 | Y | Y | Y | 97.9 | 98.9 | -2 | 2.7 | CBH and EG |
| 367 | W | W | W | 94.3 | 98.6 | -1 | 3.0 | CBH and EG |
| 171 | Y | Y | Y | 98.0 | 97.3 | -1 | 3.8 | CBH and EG |
| 216 | W | W | W | 97.9 | 68.1 | -1 | 5.6 | CBH and EG |
| 228 | H | H | H | 96.6 | 97.5 | 1 | 2.8 | CBH and EG |
| 376 | W | W | W | 96.8 | 99.1 | 2 | 3.5 | CBH and EG |
| 252 | Y | Y | - | 85.0 | 17.2 | 2 | 5.8 | CBH |
| 381 | Y | Y | - | 92.8 | 0.2 | 2 | 3.5 | CBH |

\*The positions are listed in order of proximity to the glycosyl subsites. Aromatic positions conserved in both CBHs and EGs are near the catalytic center, whereas aromatic positions conserved in only CBHs flank the catalytic center. All conserved positions are shown on the crystal structure of *TreCel7A* in **Fig. 8**.

When these positions are viewed on the crystal structure **(Table 3, Figure 8)**, an interesting pattern is observed. Whereas positions which are strongly conserved in both CBHs and EGs (first group) are located near the catalytic center of the active site, positions which are conserved in CBHs but not in EGs flank the catalytic center nearer to the “substrate-binding” sites (−7 to −1) or the “product-binding” sites (+1 to +2).

**FIGURE 8:**
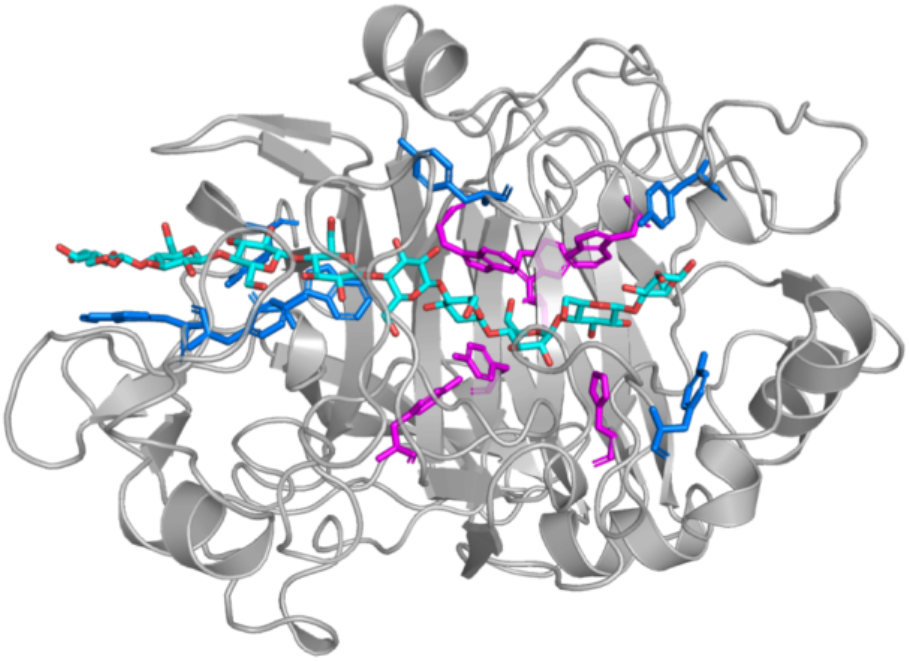
Conserved aromatic residues in the active site of *Tre*Cel7A (PDB code: 4C4C) within 6 Å of the cellononaose ligand. Residues in magenta are conserved (>66% frequency) in both GH7 CBHs and EGs and are found close to the catalytic center between −1 and +1 glycosyl subsites. Residues in blue are conserved in GH7 CBHs but not in EGs and flank the catalytic center.

### Predicting the presence of CBMs with machine learning: relationships between the CD and the CBM

The CD of GH7 proteins may be attached to a second domain (the CBM) via a flexible linker. The CBM function is mostly attributed to enhancing the binding of the enzyme to the cellulose substrate, and thus, facilitating turnover by increasing enzyme concentration on the cellulose surface (2).

We studied the distribution of family 1 CBMs in our dataset of 1,748 GH7s. First, a database of the 1,748 sequences was created and then a BLAST search of *Tre*Cel7A CBM was performed against the database. From a careful manual inspection of the BLAST alignment output, we selected an alignment score of 30 as the threshold so that GH7 sequences which yielded BLAST alignment scores of 30 or greater were determined to possess a family 1 CBM. We compared the distribution of CBMs among GH7 CBHs and EGs in our dataset and determined that 27% of GH7s contain a CBM, with 31% and 15% of GH7 CBHs and EGs exhibiting CBMs, respectively **(Table 4)**. Thus, GH7 CBHs appear to be roughly two times more likely than EGs to contain a CBM. Moreover, a chi-square test of independence indicated that the relationship between CBM utilization and GH7 subtype (CBH/EG) is significant (*p* < 0.001).

**Table 4.** Distribution of CBMs in GH7s showing the relationship between subtype (CBH/EG) and the presence of a CBM*.

|  | CBH | EG | Total |
| --- | --- | --- | --- |
| Has CBM | 407 | 66 | 473 |
| No CBM | 899 | 376 | 1275 |
| Total | 1306 | 442 | 1748 |
| CBM frequency (%) | 31.2 | 14.9 | 27.1 |

\*GH7 CBHs are roughly two times more likely to possess a CBM than GH7 EGs ( $p < 0.0001$ , chi-square test)

To investigate relationships between the CD and the CBM, we applied ML to predict the presence of CBMs using the specific amino acid residues in the CD as features. Positions flanking the CD in the MSA were removed and one-hot encoding was applied to transform the amino acids in the MSA to binary variables (72). Therefore, the MSA was transformed to a matrix such that the rows indicate the sequences, and columns denote the amino acid at positions in the MSA (features). Columns are labeled as “residue-position” and can take values of 0 or 1. For example, a value of 1 at columns Q1 and S2 for *Tre*Cel7A indicate that Gln and Ser are present at positions 1 and 2 in the MSA, respectively (**Figure 9B**). Subsequently, one-hot encoding resulted in a high-dimensional matrix with 1,748 rows and 5,933 columns. We implemented the random forest algorithm (73) with 500 trees to predict the presence of a CBM using the 5,933 one-hot encoded features. The random forest algorithm is especially suitable for this classification problem because it is capable of robustly dealing with high dimensional data by performing implicit feature selection in the learning process (74), is more tolerant to noise and overfitting (73,75), and can be used to evaluate the relative importance of the features (76).

**FIGURE 9:**
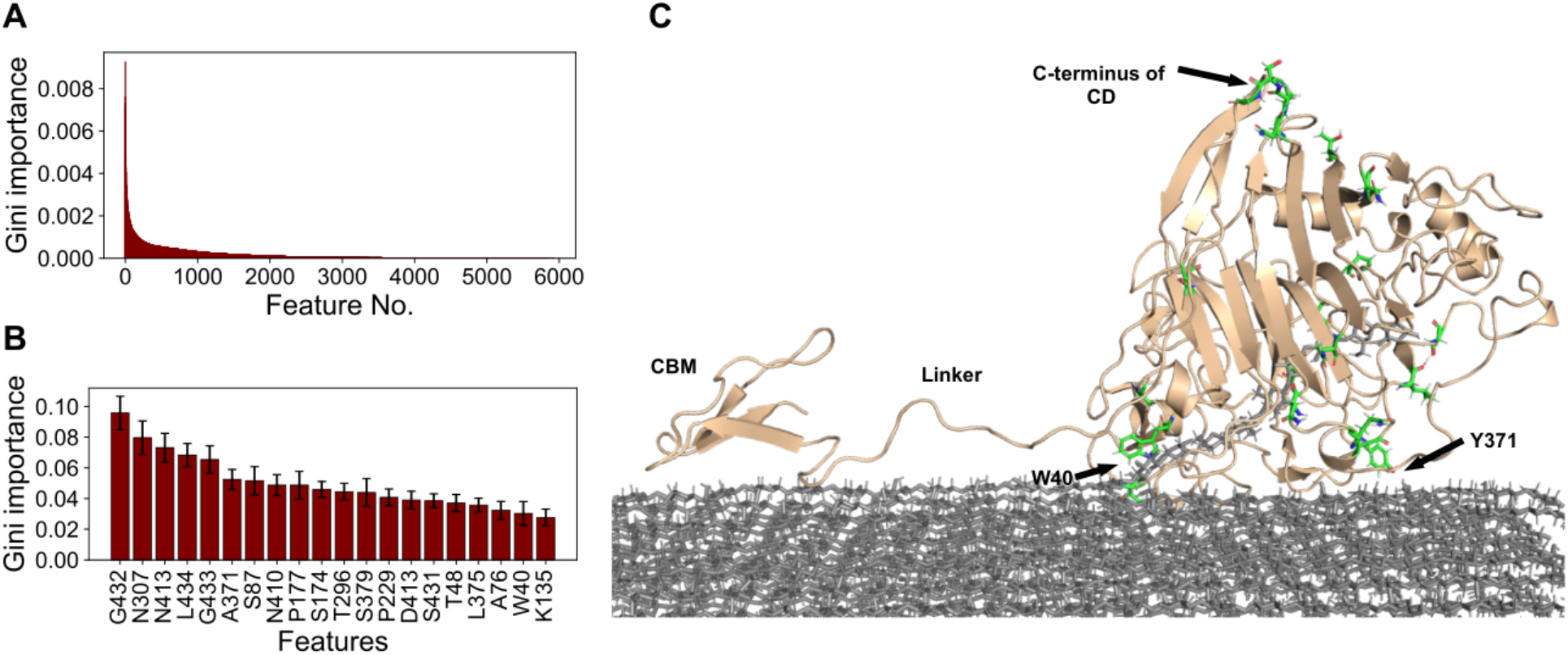
Top-performing features of the random forest classifier in predicting the presence of CBMs in GH7s. **(A)** Relative Gini importance of all 5,933 features derived from one-hot encoding of the MSA. Most features provide little information to the model **(B)** Relative (Gini) importance of top 20 features in the random forest classifier retrained on only top 20 features. Error bars indicate standard deviation measured over 100 repetitions of five-fold cross validation. **(C)** Residues of top 20 features (green sticks) shown on the structure of *Tre*Cel7A (tan cartoon) on cellulose (gray sticks). The structure is derived from a snapshot (t = 0.73 μs) of MD simulations conducted in a previous work (100).

The performance of the random forest classifier was evaluated with 100 repetitions of five-fold cross validation with random undersampling, as described previously **(Figure 4)**. Only 90% of the dataset was used for the cross validation; 10% of the dataset (174 sequences) was randomly selected and set aside for a separate final test. The random selection of the test dataset was implemented in such a way that a similar distribution (27% CBM, 73% no CBM) was maintained. In the validation routine, an accuracy of 90.8% was achieved by the 500-trees random forest trained on all 5,933 features **(Table 6)**. A plot of the relative (Gini) importances (76) of the features shows that most of the 5,933 features contribute little or no information to the performance of the random forest classifier (**Figure 9A**). We reapplied the random forest algorithm using only the top 50 and the top 20 features with the highest Gini importances. The classifiers trained on only the top 20 and top 50 features showed fairly similar validation performance to the classifier trained on all 5,933 features (**Table 6**). Some residues at the C-terminus of the CD (where the CD connects with the CBM-linker domain) were identified to be among the most important positions in predicting the presence of CBMs **(Figure 9B, Table S2)**.

To confirm that the random forest algorithm was not predicting the presence of a CBM mainly by looking at these inter-domain connecting residues (S431, G432, S433, G433, T433, L434), we repeated the validation procedure with the top 50 features but excluded features derived from positions near the C-terminus (6 features removed, 44 features remaining). The results show that the performance of the new classifier trained on 44 features was only slightly lower, with the accuracy dropping by less than three percent. Moreover, on the separate test set, the classifier trained on the top 20 features achieved an accuracy of 89.7%, confirming that the presence of a CBM can be predicted from a few residues in the catalytic domain with considerable accuracy. In addition, we derived position-specific classification rules with each of the top 50 features, as described previously (i.e. X=>CBM, else, no CBM). As expected, all 50 rules independently performed worse, compared to the random forest classifier trained on all the 50 features (MCC <0.60, vs 0.81, see **Table S2**). Among these 50 rules, the top six rules are derived from L434, G433, T433, G432 (C-terminus residues), and C4 and C72, which are the Cys residues in *Tre*Cel7A that form a rare disulfide bridge (**Table 5 and Figure S7**) (21).

**Table 5.** Distribution of CBMs in GH7s showing the relationship between the presence of the rare disulfide bond (C4-C72 in *Tre*Cel7A) and the presence of a CBM*.

|  | Has<br>disulfide<br>bond | Lacks<br>disulfide<br>bond | Total |
| --- | --- | --- | --- |
| Has CBM | 105 | 368 | 473 |
| No CBM | 54 | 1221 | 1275 |
| Total | 159 | 1589 | 1748 |
| CBM frequency (%) | 66.0 | 23.2 | 27.1 |

\*GH7s possessing this disulfide bond are roughly three times more likely to possess a CBM than GH7s lacking the disulfide bond ( $p < 0.0001$ , chi-square test)

## Discussion

In this study, we apply data mining techniques to investigate relationships between sequence and function of GH7s. We are able to accurately discriminate 1,748 GH7 CBHs and EGs with ML using only the number of residues in the active-site loops as features. However, whereas the ML models trained on the lengths of A4, B2, B3, and B4 loops achieved high predictive performance (>94% accuracy), the models trained on the other loops demonstrated mediocre or poor performance **(Table 1, Figure 5A)**. These results indicate that the lengths of the A4, B2, B3, and B4 loops are primarily important for the difference in GH7 CBH and EG behavior. Greater exposure of the active site is generally accepted as a hallmark of nonprocessive cellulases (EGs). In addition, the ML results indicate that exposure of the active site in GH7 EGs occurs primarily at the product-binding region (+1 and +2 glycosyl subsites) due to deletions in the A4 and B4 loops, at the region below the catalytic center due to deletions in the B3 loop, and at the region to the lower left of the catalytic center due to deletions in the B2 loop **(Figure 1 and 5C)**.

Earlier works have indicated that GH processivity correlates with ligand binding affinity, ligand solvation, and the flexibility of catalytic residues (33,77,78). In *Tre*Cel7A, binding affinity is stronger at product-binding sites (+1, +2) than at the substrate-binding sites (−7 to −1), and this binding affinity difference has been proposed to be the driving force for the forward processive motion of the cellulase chain (8,33,36,79). Consequently, a logical explanation for why the lengths of the A4 and B4 loops strongly correlate with GH7 CBH/EG function is as follows: deletions in the A4 and B4 loops increase ligand solvation, disrupt protein-substrate hydrogen bonds, and lower binding affinity at the product binding sites, leading to a decrease in processivity. Similarly, the strong relationship between the lengths of the B2 and B3 loops and GH7 CBH/EG function can be explained by the rationale that deletions in the B2 and B3 loops lead to an increase in solvation and a decrease in protein-ligand interactions in the substrate-binding sites, and an increase in solvation and flexibility of catalytic residues. It is interesting that although the A2 and A3 loops also overlay the catalytic center of the active site, their lengths show practically no correlation with GH7 CBH/EG function **(Figure 5A and S1),** and exposure of the catalytic center in GH7s is achieved primarily by deletions in the B2 and B3 loops instead.

Moreover, the level of variation in lengths of the loops, as measured by the relative standard deviation, positively correlates with the predictive performance of the loops in discriminating GH7 CBHs and EGs **(Figure 5A and 5B)**. This suggests that variation in the lengths of active-site loops was a major strategy in the evolutionary design of processivity in GH7s so that variation was allowed in the loops that significantly affect processivity and limited in other loops that have little impact on processivity (A2 and A3).

Furthermore, there is a strong positive correlation between the lengths of the A4, B2, B3, and B4 loops **(Figure 6)**. Hence, in wild type GH7s, the shortening of any one of these four loops is highly associated with truncation of the other three loops. On our dataset of 1,748 sequences, we observed that if the B4 loop of a sequence is shortened, as is typical of GH7 EGs (i.e. possessing three residues or less), the probability that the A4, B2, and B3 loops are all shortened to typical GH7 EG lengths (i.e. five, four, and three residues or less, respectively) is 0.97 **(Figure 5D)**. In other words, the pronounced concurrent shortening of the A4, B2, B3, and B4 loops observed in all crystal structures of GH7 EGs (27,30,80) is remarkably conserved in EGs across the GH7 family. This distinct bimodal distribution (**Figure S1**) and strong correlation between loop lengths and GH7 subtype may serve as a valuable tool for correct gene annotation. Moreover, the strong conservation of loop lengths also indicate that there are coupled interactions between the A4, B2, B3, and B4 loops (81). This might explain why in the recent work of Schiano-di-Cola *et al*., independent deletions in the B3 and B4 loops did not lead to significant improvements in the activity of *Tre*Cel7A on amorphous cellulose, and deletions in the A4 loop rendered the enzyme inactive (53). Cooperative synergy of deletions in other loops, as well as point mutations at key positions, may be required to fully exploit the effects of deletions in the B3, B4, and A4 loops.

Beyond the active-site loops, we have derived 90 position-specific classification rules from 42 positions in the MSA, such that the specific amino acid, or amino acid type, at any of these positions can independently predict the subtype of GH7s, with accuracies ranging from 87% to 97%. The high accuracy of these classification rules implies that there are strong constraints on the specific amino acids, or amino acid types, utilized by GH7 CBHs and EGs at these 42 positions. Such differential constraints likely signify that these positions play imperative roles in the difference between GH7 CBH and EG behavior. More than half of these positions are within 5 Å of the cellononaose substrate bound in the *Tre*Cel7A structure, and many of these positions cluster around the B2 loop (**Figure 1, 9D and 9E)**. This finding provides a possible explanation for the observation that deletions in the B2 loop led to much greater changes in the CBH-behavior of *Tre*Cel7A than deletions in the B3 and B4 loops, relative to *Tre*Cel7B (53). Since more of the important residues that yield high-accuracy position-specific classification rules cluster around the B2 loop than other loops, deletions in the B2 loop likely leads to a disruption of a greater number of interactions necessary for CBH activity than deletions in the B3 and B4 loops.

Many of the 42 positions from which we derived classification rules have been identified and studied in previous works, and mutations at these positions have led to significant increase in catalytic efficiency (82–84). Trp38, Tyr51, Asn103, Lys181, Asn200, Asp179, Arg251, Asp262, and Arg394 were identified in a docking study as residues that directly interact with and stabilize the cellulose substrate in the active site of *Tre*Cel7A (85). Several of these positions have been further shown to form important stabilizing interactions with the substrate. Arg394 forms hydrogen bonds with the +2 glycosyl residue (35,36), Arg251 forms a salt bridge with Asp259 and hydrogen bonds with the +1 and +2 glycosyl residues (6,36,37), and Asn103 and Lys181 form hydrogen bonds with the −5 glycosyl residue (19,86). Sørensen *et al*. studied mutants of *Rasamsonia emersonii* Cel7A in which two Asn residues on the B2 loop, Asn194 and Asn197 (Asn197 and Asn200 in *Tre*Cel7A, respectively) were replaced with Ala (83). They observed that the mutations led to a decrease in substrate affinity and processivity, thus, enabling faster enzyme-substrate dissociation and a corresponding increase in activity on crystalline cellulose. In this present work, the Asn200 position yields the following classification rule: Asn implies CBH, and not Asn implies EG, which discriminates GH7 CBHs and EGs with an accuracy of 89%. Similarly, Bu *et al*. conducted computational studies of several *Tre*Cel7A residues including Arg251, Asp262, and Tyr381 (37). These residues were identified to substantially interact with the cellobiose substrate and mutation to Ala resulted in considerably weaker binding of cellobiose in the product-binding site. It was suggested that these mutants would demonstrate improved biomass conversion efficiency due to accelerated expulsion of the cellobiose product. In this present study, these positions (251, 262, and 381) also yield high-accuracy classification rules with accuracies of at least 88%. Additionally, Mitsuzawa *et al*. determined that mutation of Asn63 and Lys203 to Ala in *Talaromyces cellulolyticus* Cel7A (Asn37 and Lys181 in TreCel7A, with classification accuracies of 91% and 94%, respectively) led to a remarkable increase in activity on cellulose (82).

Some positions farther away from the active site also yielded high-accuracy classification rules. For example, position 401 – conserved as Ser in CBHs but as Glu in EGs and more than 13 Å away from the cellodextrin ligand in the *Tre*Cel7A structure – generates a CBH/EG classification rule with an accuracy of 92%. Although residues at positions such as 401 may not directly interact with the cellulose substrate in the active site, they may participate in long-range interactions that affect GH7 CBH and EG behavior. Further studies are required to determine the specific roles these conserved positions play in function and structural stability. Altogether, we surmise that the positions that yield high-accuracy classification rules play key roles in GH7 CBH/EG function and, as such, should be carefully considered when engineering the protein at or around these sites.

Bioinformatic analysis of the MSA revealed conserved aromatic positions in the active site that are within 6 Å of the cellulose substrate in *Tre*Cel7A **(Table 3, Figure 8)**. The results indicate that whereas conserved aromatic residues in the active site of GH7 CBHs span the entire active-site tunnel, conserved aromatic residues in the active site of GH7 EGs are clustered around the catalytic center. Moreover, aromatic positions near the catalytic center are conserved in both GH7 CBHs and EGs. This arrangement of conserved aromatic residues in the active site suggests that while aromatic residues near the catalytic center (Y145, W216, H228, W367, and W376) play major roles in catalytic bond cleavage, conserved aromatic residues that flank the catalytic center (W38, W40, Y51, Y82, Y252, Y370, and Y381) are utilized mainly by CBHs for processive motion. Several experimental and computational studies support this hypothesis (37,39,87,88).

Taylor *et al*. assayed chimeras derived from interchanged subdomains of *PfuCel7A* and *Tre*Cel7A. Although the CD of *PfuCel7A* exhibited greater efficiency on biomass than the CD of *Tre*Cel7A, interchanging CBM and linker regions did not yield a uniform trend in catalytic efficiency. As a result, it was concluded that there are complex interactions that are not yet well-understood between the domains (21). In this work, we have applied ML to predict the presence of CBMs from amino acid positions in the CD to map relationships between the CBM and CD of GH7s. First, our data indicate that GH7 CBHs are roughly two times more likely to utilize CBMs than GH7 EGs, which is as expected since CBMs likely enable CBHs stay longer on the cellulose substrate to facilitate consecutive hydrolysis. Furthermore, ML results show that the presence of a CBM in GH7s can be accurately predicted (89.3%) using only 20 features derived from 19 positions in the catalytic domain **(Table 6)**. This high predictive accuracy largely suggests that there are constraints and key functional relationships between residue positions in the CD and the presence of a CBM in the gene. Interestingly, on the protein structure, these 19 positions are mostly located on loops or at turns all over the protein structure **(Figure 9C)**. Moreover, the amino acid residues constituting the 20 features are mostly small amino acids (such as Gly, Ser, Thr, Asp, and Asn) that are known to affect the conformational flexibility of proteins (89,90). Taken together, our ML results, while preliminary, suggest that the presence of CBMs in GH7s correlates with the overall conformational flexibility of the CD, and that CBMs may exist, in part, to compensate for highly flexible CDs that are more likely to detach from the cellulose surface. Moreover, the position-specific rules we derived from the top 50 random-forest features in predicting CBMs indicate that GH7s possessing a rare disulfide bond (C4-C72 in *Tre*Cel7A) are about three times more likely to possess a CBM than GH7s that lack this disulfide (**Table 6**, **Table S2, Figure S7**). In a previous work, mutation of C4 and C72 in *Tre*Cel7A was shown to increase cellulolytic efficiency and flexibility of the tunnel entrance (21). Since an extra disulfide bridge would generally decrease the flexibility of the CD, the correlation of C4 and C72 with the presence of a CBM is contrary to our hypothesis that CBMs compensate for the flexibility of the CD. This paradoxical correlation, thus, warrants further experiments to investigate such relationships.

**Table 6.** Performance (%) of random forest classifiers in predicting presence of CBM*.

|  | Validation |  |  |  | Testing |
| --- | --- | --- | --- | --- | --- |
|  | All 5,933 features | Top 50 features | 44 features (no C-terminus) | Top 20 features | Top 20 features |
| Accuracy | 90.8 ± 2.1 | 90.9 ± 2.1 | 88.2 ± 2.5 | 89.3 ± 2.4 | 89.7 |
| Sensitivity | 93.7 ± 2.8 | 92.2 ± 2.9 | 89.6 ± 3.4 | 90.0 ± 3.2 | 95.7 |
| Specificity | 87.9 ± 3.5 | 89.7 ± 3.3 | 86.9 ± 3.7 | 88.5 ± 3.6 | 87.4 |
| MCC | 0.80 ± 0.05 | 0.81 ± 0.05 | 0.76 ± 0.05 | 0.78 ± 0.05 | 0.68 |

\*Validation and testing are performed on a 90%:10% split of the dataset, respectively. Validation performance is reported as the mean over 100 repetitions of five-fold cross validation ± one standard deviation.

In conclusion, we have used ML to uncover key positions in GH7 sequences that appear to be related to function and statistical relationships between GH7 sequence and functional diversity. While these relationships are statistically significant, we stress that they may be influenced by sampling and phylogenetic biases inherent to the dataset. Nonetheless, as the ML strategies we have applied to GH7s may be extended to other protein families, particularly where multiple functional classes exist in the family (such as CBH/EG or CBM/no-CBM), this work provides a solid basis for the statistical investigation of sequence-function relationships in protein families. We also anticipate that the findings in this work will inform further propitious studies for the design of more efficient cellulases.

## Experimental procedures

### Sequence datasets

Sequences were retrieved by protein-protein BLAST searches against the NCBI non-redundant database by using *Tre*Cel7A (P62694.1) and *FoxCel7B* (AAA65586.1) as query sequences. BLAST search was implemented with the NCBI web server (https://blast.ncbi.nlm.nih.gov/Blast.cgi) using default settings. Only sequences with E-values of 1e-20 or better and query cover of 60% or more were retained. The query cover threshold of 60% was applied to exclude the large number of fragment sequences returned by the BLAST search. A total of 2,024 sequences were retrieved. A sequence identity threshold of 99% was applied to remove redundant sequences so that only 1,748 sequences were left in the dataset. From manual inspection of the BLAST output, 60 of these sequences consisted of multiple domains other than GH7. Other domains were deleted in these sequences leaving only one GH7 domain for each sequence. The SwissProt/UniProt dataset of 44 sequences was obtained by a similar BLAST search against the SwissProt/UniProt database.

### Sequence alignments

Sequence alignments of the SwissProt/UniProt dataset (44 sequences) and the annotated NCBI dataset (427 sequences) were conducted with MAFFT version 7 (91) using BioPython (92) with default settings. Due to the greater diversity of the larger dataset (1,748 sequences), in order to avoid generating erroneous alignments, a structure-based sequence alignment was implemented for the larger dataset. First, structural alignment of 20 GH7 structures (16 CBHs, 4 EGs) was conducted with the Promals3D web server (93). The structural alignment was manually edited in UGENE (94) following standard manual adjustment methods (95). Then, an MSA of the 1,748 sequences was generated with the MAFFT add-sequences option (91) by adding the sequences to the structural alignment.

Sequence alignments were viewed with ESPript (http://espript.ibcp.fr) (96), and sequence logos **(Figure S4)** were generated with WebLogo (https://weblogo.berkeley.edu/logo.cgi) (97).

### Machine learning and performance evaluation

Profile hidden Markov models were constructed from the MSAs with a local version of the HMMER software (version 3.1b2) (57,98). All ML methods were implemented using the Scikit-learn Python package (version 0.20.3) (99). The K-nearest neighbor (KNN) classifier was trained with the “n_estimators” parameter (k) set to an optimal value of 10 (best of 5, 10, and 15). A radial basis function (RBF) kernel was applied in the support vector machine (SVM) classifiers, and default settings were used for the logistic regression classifiers. To avoid overfitting with the decision trees, the depth of the trees was limited to the number of features. Hence, single-feature decision trees had a “max-depth” of one, and the decision tree trained on all eight features had a “max-depth” of eight.

There were severe outliers in the lengths of active-site loops that would have skewed the ML results. For example, from the MSA, a sequence (GenBank accession: CRK24563.1) had 140 residues in the B2 loop. These extremities may have resulted from sequencing or splicing errors. Before the ML procedure, outliers were capped to an arbitrarily selected maximum limit (62) **(Figure S5)**. All nonbinary features applied in ML were standardized by converting them to Z-scores according to following equation:

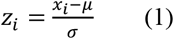

The ML algorithms were applied to discriminate between a positive class (CBH or CBM) and a negative class (EG or no CBM), resulting in four classification outcomes: true positives (TP), true negatives (TN), false positives (FP), and false negatives (FN). The performance of the ML algorithms was evaluated by computing the sensitivity, specificity, accuracy, and MCC according to the following equations:

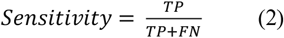

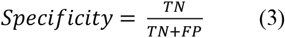

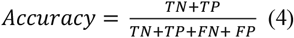

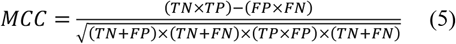

## Data availability

All datasets and Python scripts used in this study are available at https://doi.org/10.5281/zenodo.4216573.

## Acknowledgements and funding information

This work was supported in part by the National Science Foundation (CBET-1552355 to CMP in support of JEG). This work was also authored in part by the Alliance for Sustainable Energy, LLC, the manager and operator of the National Renewable Energy Laboratory for the U.S. Department of Energy (DOE) under Contract No. DE-AC36-08GO28308. Funding was provided to GTB by the U.S. Department of Energy Office of Energy Efficiency and Renewable Energy Bioenergy Technologies Office. This material is also based upon work supported by (while CMP is serving at) the NSF. Any opinion, findings, and conclusions or recommendations expressed in this material are those of the authors and do not necessarily reflect the views of the NSF.

